# A sequence-based classifier distinguishes phenotype-associated genes from other gene models in plants

**DOI:** 10.64898/2025.11.30.691407

**Authors:** Nikee Shrestha, Zhongjie Ji, Xiuru Dai, Pinghua Li, James C. Schnable

## Abstract

Only a small fraction of annotated plant genes possesses experimentally validated associations with specific phenotypes. Phenotype associated genes have distinct structural, molecular, and evolutionary characteristics compared to non-validated gene models. Here, we developed a simple classifier that uses sequence and evolutionary features which can be generated for any species with an annotated reference genome assembly, to accurately distinguish phenotype associated genes from both the overall population of annotated gene models and a specific set of genes identified as being tolerant of loss of function mutations. A model trained solely on genes from maize (*Zea mays*) identified and prioritized rice (*Oryza sativa*) and arabidopsis (*Arabidopsis thaliana*) genes that were highly enriched in genes with experimentally validated links to phenotypes in both of these evolutionarily distant species. Gene models predicted to have a higher probability of being linked to phenotypes displayed patterns consistent with known biological properties of phenotype associated genes. Notably, the sets of genes predicted to have a high probability of being linked to phenotype variation did not consist exclusively of well-characterized gene families, but included many uncharacterized gene families carrying domains of unknown function. The quantitative scores generated by this model offer a valuable resource for prioritizing and exploring the vast number of uncharacterized gene models in plants, reducing the risk of failure in future reverse genetics efforts and potentially accelerating gene discovery and functional annotation in crops.

**Significance Statement:** Annotation of an organism’s genome produces tens of thousands of predicted genes, termed gene models. However, only a few of these have been associated with phenotypes. We built a simple algorithm that uses DNA sequences and evolutionary conservation-based information to predict which genes are most likely to change a plant’s phenotype when disrupted. In maize, it separates known phenotype-associated genes from other gene models; it also works in rice and arabidopsis. High-scoring genes show signals of biological importance and include many with no known function. This algorithm provides a shortlist of candidate for gene characterization and crop improvement.

## INTRODUCTION

The genomes of typical multicellular eukaryotes contain tens of thousands of gene model annotations. However, few of these genes have been the subject of specific genetic or molecular investigations. Indeed, in arabidopsis less than 10% of genes have been linked to specific phenotypes (Lloyd and Meinke, 2012); likewise, in rice approximately 10% of gene models are associated with phenotypes (Huang et al., 2022), and in maize this is less than 1% (Schnable and Freeling, 2011). The situation in other plants is even worse (Oellrich et al., 2015). This is not unique to plants; even in the human genome many annotated genes have not been the subject of detailed investigation by even a single study (Sinha et al., 2000), including hundreds of genes known to be essential for cell survival (Wang et al., 2015). Research continues to focus on the same well-studied genes (Kustatscher et al., 2022a,b), thereby hindering our understanding of the core functionalities of eukaryotes and the mechanisms responsible for lineage-specific processes and phenotypes.

The process of functionally characterizing less-studied gene models or generating new hypotheses about gene-phenotype associations is time-intensive and high-risk. These efforts can take years and frequently yield inconclusive or null results, creating a very real career risk for researchers. These limitations constrain both reverse genetics approaches and strategies that leverage natural genetic diversity. Marker trait association signals that fall near already known genes tend to be viewed as confirmatory. In contrast, quantitative trait nucleotides located next to gene models not previously implicated in a phenotype are less pursued, largely because functional validation requires substantial time, resources, and carries a high risk of yielding null results. Similar obstacles arise when interpreting results from differential gene expression analyses, co-expression networks, and other multi-omics approaches. In contrast, once a gene has been shown to influence at least one phenotype, the risk of investigation declines, further amplifying attention toward genes with existing functional evidence while discouraging exploratory work on the vast number of uncharacterized gene models (Kwon et al., 2024). For example, the maize teosinte branched1 (tb1) gene plays an important role in reducing tillering in domesticated maize (Doebley et al., 1997, 1990). The association of tb1 with important domestication phenotypes in maize drove the study of this gene to understand its biological role in maize and in other species (Igartua et al., 2020). As of November 2025, at least 140 publications have studied the tb1 gene (Woodhouse et al., 2012). Being able to predict, in advance, which genes are likely to influence phenotypes would reduce the risk of investigating currently uncharacterized or under characterized genes.

Genes already associated with phenotypes differ from other gene models in a range of characteristics and these properties were consistent regardless of whether the phenotype the gene was associated with was lethal, constitutive, conditional, or observable only at a biochemical or molecular level (Schnable, 2020). This implies the existence of a separate class of gene models which are either truly unrelated to determining the phenotype of an organism, or at a minimum, whose role is unlikely to be identifiable via current research approaches. Studying this class of genes is challenging. The set of gene models lacking a known phenotype comprise a mixture of both genes which could be linked to phenotype if studied and those truly unrelated to determining the observable phenotypes of an organism. The lack of incentives to publish null results makes it difficult to specifically identify the subset of gene models where researchers have looked for links to phenotypes and found none.

Here we identify a set of phenotypes-associated genes via a literature search and a set of loss-of-function tolerant genes via analysis of population scale resequencing data. We demonstrate: (i) these sets of genes are distinguished by a wide range of features, (ii) that a simple random forest classifier can predict the likelihood that an unannotated gene will be associated with a phenotype using additional phenotype-associated genes published after our initial dataset was assembled and (iii) models trained in one species can distinguish phenotype-associated genes from average gene models in closely and distantly related species without the need for retraining.

## Results

We identified 272 maize gene models linked to phenotypic outcomes based on evidence from loss-of-function alleles and named this set as phenotype-associated genes (PAGs) (Supplemental Dataset S1). Using whole-genome resequencing data from 1,515 maize genotypes, we also identified a second set of 713 gene models with variants causing a premature stop codon, i.e., loss-of-function-tolerant genes (LFTs). There was no overlap between the set of PAGs and the set of LFTs (E = 5 gene models, *p*-value = 0.017; Fisher’s exact test). This result is consistent with the hypothesis that loss-of-function alleles with obvious phenotypic consequences will be targets of stronger purifying selection than loss-of-function alleles that are silent or have only subtle phenotypic consequences. In many cases the alleles that produced premature stop codons were relatively rare, but in 162 cases, the frequency of the allele that resulted in a premature stop codon was greater than 0.25 across the resequenced population used to discover these variants (Figure 1A). We focused on these 162 genes that carry common loss-of-function variants in downstream analyses.

**Figure 1.**
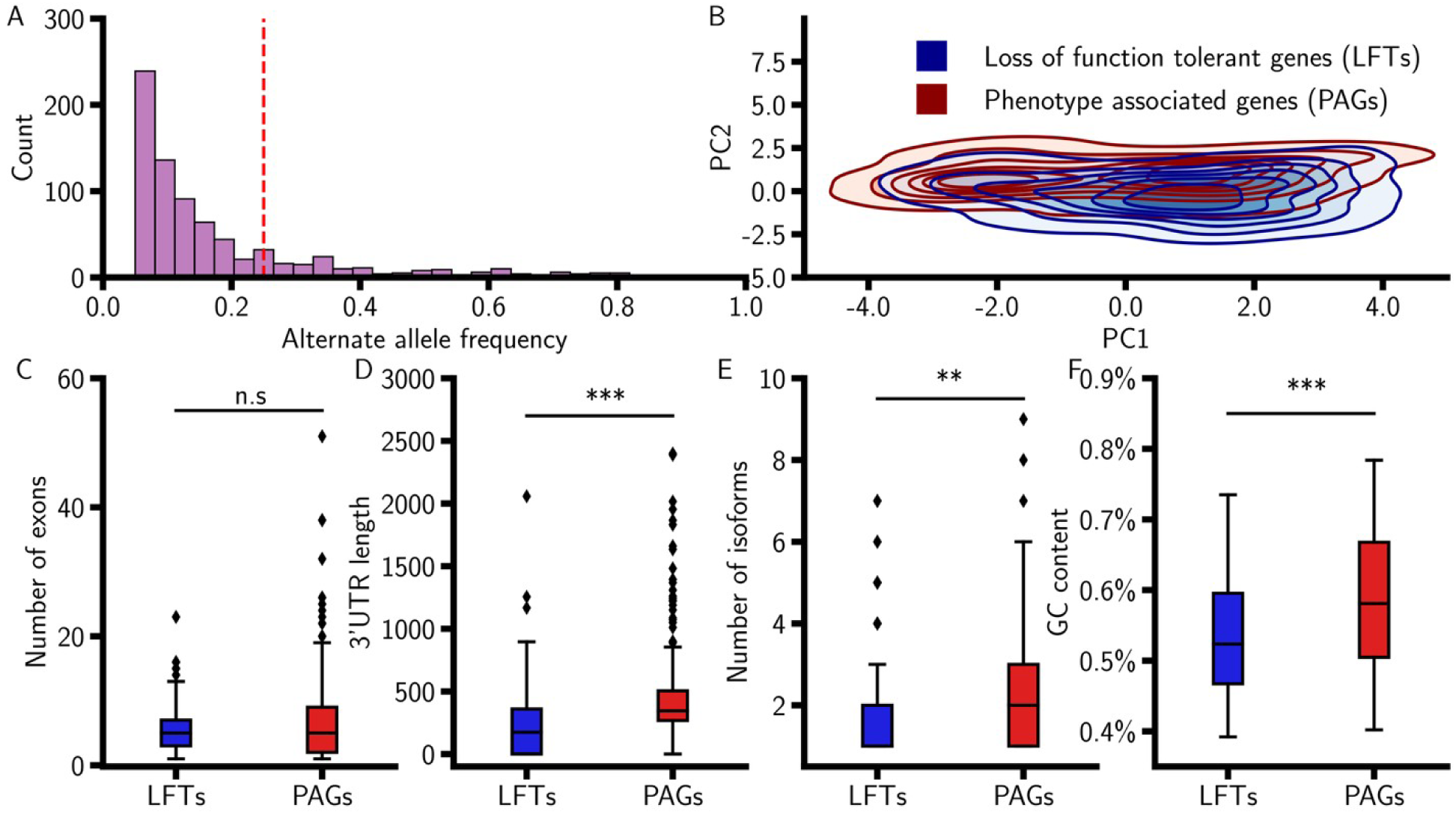
Differences between loss-of-function tolerant genes (LFTs) and phenotype-associated genes (PAGs). A). Gene models in the B73 reference genome contain alternate alleles that introduce premature stop codons, totaling 713 models. A threshold of 0.25 alternate allele frequency defined 162 loss-of-function tolerant genes (LFTs), which were used as true negatives in this study. B). Principal component analysis (PCA) of LFTs (*n*=162) and PAGs (*n*=272). The first two principal components explain 28.62% of the variance. Data included in the PCA included a set of 2,675 previously published features that were downloaded from MaizeGDB (Sen et al., 2023). C). Difference in the number of exons per gene model between LFTs and PAGs. D). Difference in the length of the three prime untranslated regions (3′UTRs) of gene models between LFTs and PAGs. E). Difference in the number of isoforms per gene model between LFTs and PAGs. F). Difference in the proportion of GC content between LFTs and PAGs. ** *P* < 0.01, *** *P* < 0.001, n.s non-significant (two-tailed Mann-Whitney U test).

Across a set of 2,604 quantitative features describing maize gene models, PAGs, and LFTs with common loss-of-function alleles exhibited statistically significant differences ∼44% of the time (*p*-value < 0.05, two-tailed Mann-Whitney U test, Supplemental Dataset S2). PAGs and LFTs exhibited significant differences in gene length, average protein abundance, the number of tissues expressing a gene model, syntenic conservation in a related species, and evidence of translation. These results are consistent with previously reported differences between genes associated with mutant phenotypes and the overall set of annotated gene models (Schnable, 2020). No significant difference was observed between the number of exons in PAGs and LFTs, another trait previously reported to differ between genes associated with mutant phenotypes and other annotated gene models (*p*-value = 0.53, two-tailed Mann-Whitney U test, Figure 1C). We speculated that defining LFTs as gene models containing premature stop codons may have introduced a bias toward longer transcripts with more exons, as longer coding sequences provide more opportunities for mutations to produce a premature stop codon. As a result, the LFTs may be enriched for longer transcripts with more exons, preventing any true difference in the number of exons between LFTs and PAGs. Many other features where PAGs and LFTs significantly differed had not previously been linked to the likelihood of being associated with organismal phenotypes. These include the length of the 3’ untranslated region (UTR) (PAGs > LFTs *p*−value < 0.001 [8.13e-15], two-tailed Mann-Whitney U test, Cohen’s d=0.64,Figure 1D), number of isoforms (PAGs >LFTs *p*−value= 0.003, two-tailed Mann-Whitney U test, Cohen’s d=0.23, Figure 1E), and GC content in the nucleotide sequence of the gene model (PAGs >LFTs *p*−value < 0.001 [6.70e-5], two-tailed Mann-Whitney U test, Cohen’s d=0.53, Figure 1F). PAGs appeared to be more distinct from the overall population of annotated gene models other than LFTs, with significant differences between PAGs (mean Cohen’s d= 0.05, median Cohen’s d= 0.01) and non-PAG non-LFT genes for 54% of quantitative features and significant differences between LFTs (mean Cohen’s d= 0.008, median Cohen’s d= 0.002) and non-PAG non-LFT genes were observed for only 28% of quantitative features (Supplemental Dataset S2). Collectively, these results demonstrate that LFTs are more similar to the rest of the gene models than PAGs, and that PAGs are significantly different from both LFTs and the rest of the average gene models for both previously tested and untested features.

Despite the significant differences between PAGs and LFTs for many individual features, these two populations of genes exhibited only modest separation on the first principal component calculated from the full set of gene features (Figure 1B). This result contrasts with the previously reported clear separation between phenotype-associated genes and other annotated gene models when using a smaller, curated feature set (Schnable, 2020). A random forest model trained to distinguish PAGs and LFTs using all 2,675 gene features exhibited strong predictive performance, achieving an ROC curve with an AUC of 0.89 on the hold-out test gene sets (Figure 2A). The prediction probabilities assigned to PAGs (mean=0.86) and LFTs (mean=0.23) were each significantly different from those of uncharacterized annotated gene models (mean=0.50, *p* − value < 0.001 for both comparisons, two-tailed Mann-Whitney U test, Supplemental Dataset S3). Thirty percent of all uncharacterized annotated genes in the maize genome were classified as more likely to be loss-of-function tolerant than the lowest-scoring PAG in the dataset (Figure 2B). The ability of a random forest model to achieve strong predictive performance on hold-out test sets of LFTs and PAGs supports the substantial differentiation of these populations of gene models.

**Figure 2.**
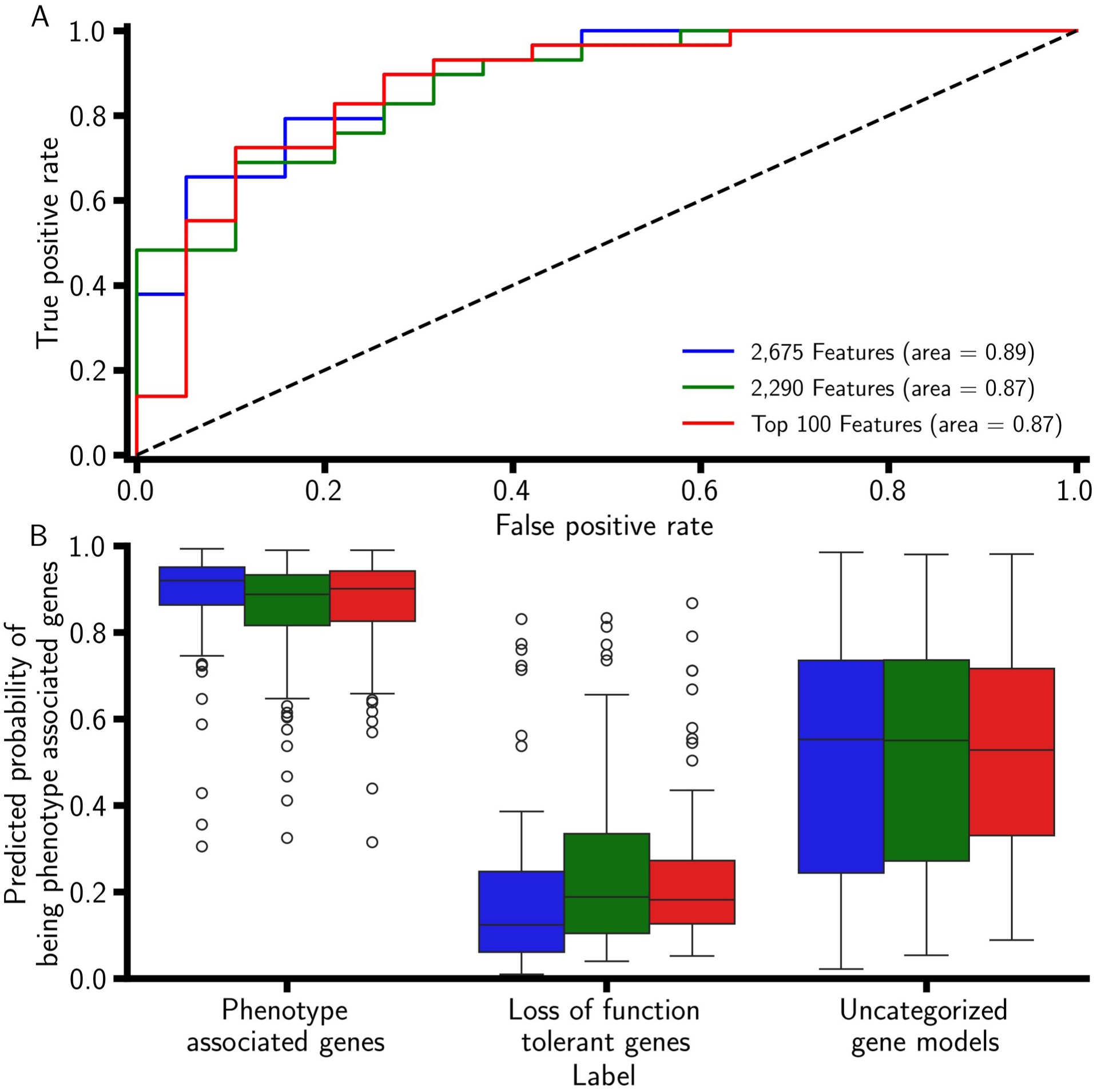
Model performance using different subsets of total features. A). ROC curves are generated from three models: one using all 2,675 features (experimental and genomic sequence-derived), one using 2,290 non-experimentally derived features, and one using the top 100 non-experimental sequence-derived features. Performance was evaluated on 48 withheld gene models. B). Predicted probability from the three models in Panel A across all maize gene models.

We next sought to determine how many, and which types of features were necessary to distinguish these populations of genes. The set of features describing maize gene models can be separated into sequence-derived and experimentally derived features. Sequence-derived features can be directly extracted or calculated, given only an assembled genome sequence and a structural gene annotation file. By contrast, experimentally derived features, including gene expression data, protein abundance, chromatin accessibility profiling, and assays of transcription factor binding, require additional experiments and are typically challenging to generate similarly across experiments conducted in multiple species. Among the 100 features with the highest Gini importance in the trained PAG/LFT classification model, 21 were experimentally derived; however, none of the 10 features with the highest Gini importance were experimentally derived (Supplemental Dataset S4). A random forest model trained solely on non-experimentally derived features achieved comparable performance, with an ROC curve area of 0.87 (Figure 2A, B). Model performance remained consistent when the feature set was restricted to only 100 non-experimentally derived features with the highest Gini importance (Figure 2A, B). The probability of a gene in the top 10% of predicted scores generated by the hundred-feature no-experimental data model being linked to a phenotype was estimated to be approximately 22.5 times higher than that of a gene with a score in the bottom 50% based on evaluation of the hold-out test data. These results suggest that it may be feasible to build high-performing models based solely on feature sets that can be generated consistently across different species.

The genetic characterization of several plant species, notably rice and arabidopsis, also includes information on the phenotypic consequences of disrupting comparable or greater numbers of genes compared to the maize dataset used in this study. However, in most plant species, with sequenced genomes, few or no genes have been directly linked to mutant phenotypes. We assessed the transferability of a model trained on maize data to other species using a model trained on 95 of the top 100 maize features, excluding five that could not be easily generated for rice gene models (see Methods), to predict the probability that annotated rice genes would be associated with phenotypic consequences. A set of 94 rice PAGs was assigned significantly higher probabilities of being associated with a phenotype than with the background set of annotated gene models in the rice genome (Figure 3A, *p*−value < 0.001 [2.74e-14], two-tailed Mann-Whitney U test, Supplemental Dataset S5). The probability of a rice gene in the top 10% of scores being linked to a phenotype was six times higher than that of a gene in the bottom 50%. The divergence of the lineages leading to maize and rice is estimated to have occurred approximately 50 million years ago, roughly one-third the time since the estimated age of the most recent common ancestor of maize and arabidopsis (∼160 million years ago) (Kumar et al., 2017). The arabidopsis genome is also less than 10% the size of the maize genome, with substantially lower transposon content. A model trained solely on maize data using the same 95 features assigned significantly higher probabilities of being associated with a phenotype to a set of arabidopsis PAGs than to the background set of annotated gene models in the arabidopsis genome (Figure 3B, *p*−value < 0.001 [7.86e-108], two-tailed Mann-Whitney U test, Supplemental Dataset S5). The probability of an arabidopsis gene model in the top 10% of scores being linked to a phenotype was more than three times higher than a gene in the bottom 50%. These results confirm the notion that a model trained solely on maize data can still achieve both statistically significant and operationally meaningful performance in prioritizing genes for phenotypic characterization when applied to a distantly related angiosperm with a radically different genome architecture.

**Figure 3.**
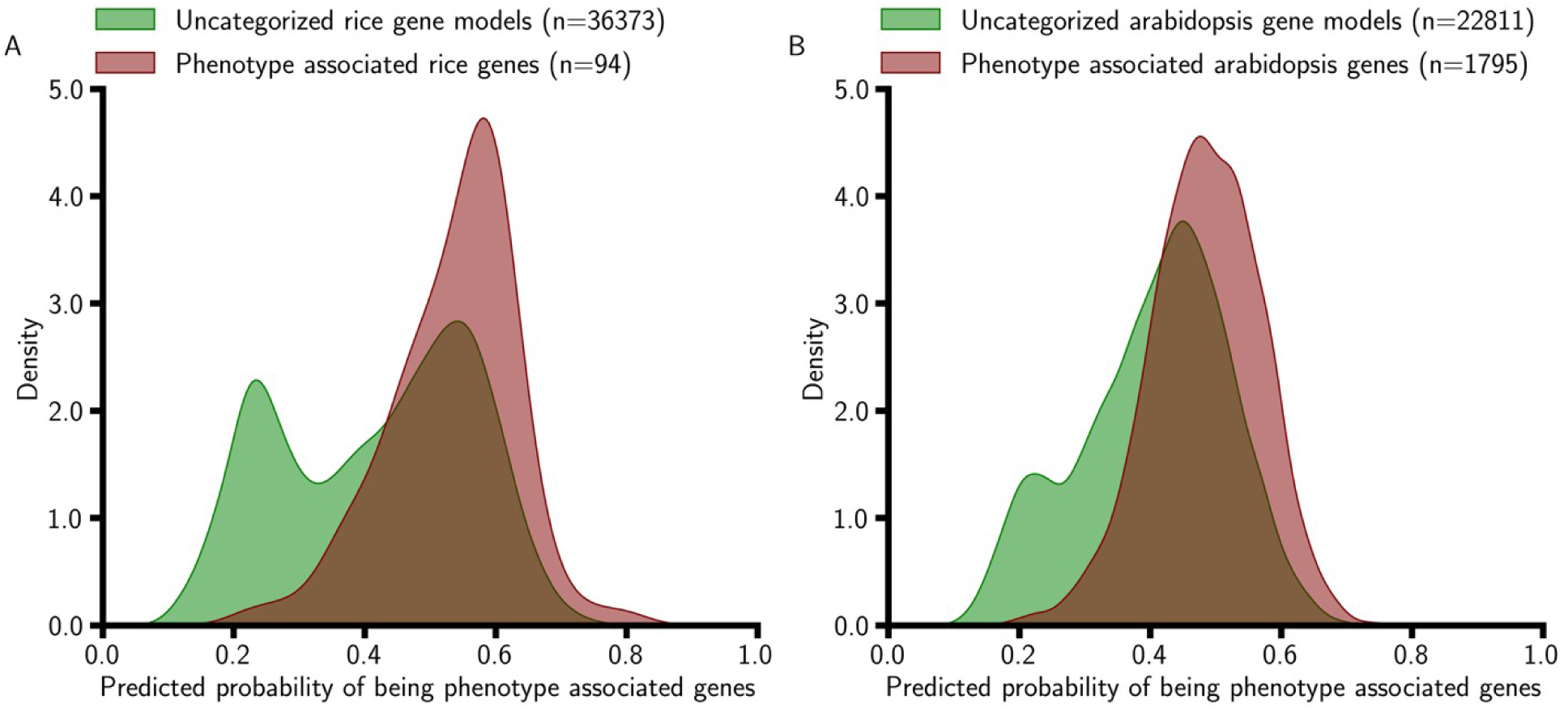
Phenotype-associated genes have higher predicted probabilities than uncategorized gene models in rice and arabidopsis. A). Distribution of predicted probabilities of a subset of validated gene models (*n*=94) retrieved from the literature in rice (Huang et al., 2022) using a model trained on maize data with 95 non-experimentally derived features. B). Distribution of predicted probabilities of a subset of validated gene models (*n*=1,922) retrieved from the literature in arabidopsis (Lloyd and Meinke, 2012) using a model trained on maize data with 95 non-experimentally derived features.

The gene models predicted to be linked to phenotypes are supported by other data types associated with purifying selection or the consequences of mutations. Gene models in the bottom quartile of probability of being associated with a phenotype exhibited approximately 4.6 times higher odds of containing a premature stop codon compared to those in the top quartile (*p*-value < 2.2e-16; Fisher’s exact test) (Figure 4A, B). Variants introducing premature stop codons in bottom-quartile gene models were observed at higher minor allele frequencies than variants introducing premature stop codons in top-quartile genes (*p*-value = 0.002, two-tailed Mann-Whitney U test). Variant frequency per kilobase of coding sequence (CDS) was higher in top-quartile gene models, but this was explained entirely by a higher prevalence of variants predicted to have small (low) effects on protein function, typically synonymous variants, and the proportion of variants predicted to have moderate (*p*−value < 0.001, two-tailed Mann-Whitney U test) or strong (high) effects on protein function (*p*−value < 0.001, two-tailed Mann-Whitney U test) were significantly higher in bottom-quartile gene models (Figure 4C, D). Not all functional consequences of DNA substitutions will be reflected in estimates of the effects of these substitutions on protein function. This gap can be addressed in part using DNA language models, which can use context information to estimate the effects of DNA substitutions. Genetic variants in the gene body, mRNA, or CDS of top-quartile gene models exhibited significantly higher predicted likelihoods of being deleterious (based on average absolute zero-shot scores from PlantCaduceus) than bottom-quartile gene models (*p*−value < 1e-200, two-tailed Mann-Whitney U test, Figure 4E). By contrast, variants present in UTRs or introns exhibited a small but significant bias in the opposite direction, with variants in these regions of bottom-quartile genes assigned marginally higher absolute zero-shot scores by PlantCaduceus than those in top-quartile genes (*p*−value < 1e-20, two-tailed Mann-Whitney U test, Figure 4E). Constraining GO enrichment analysis to the population of only those genes assigned at least one GO term, out of 7,277 GO terms assigned to at least one gene model, top-quartile genes were enriched in 891 GO terms and bottom-quartile genes were enriched in only 30 GO terms (Supplemental Dataset S6). This result raised the concern that the model’s strong performance might reflect memorization of the characteristics of well-annotated gene families. Ninety-six percent of top-quartile genes and forty-seven percent of bottom-quartile genes were assigned at least one GO term. However, 162 genes in the top decile of predicted probabilities of being phenotype-associated encode proteins with domains of unknown function. Manual curation of the top 15 gene models showed that more than half had only domains of unknown function. These results indicate that the model captured biologically important patterns to distinguish gene models highly likely to be associated with phenotype, and that genes predicted to be associated with phenotypes extend beyond those of well-characterized gene families.

**Figure 4.**
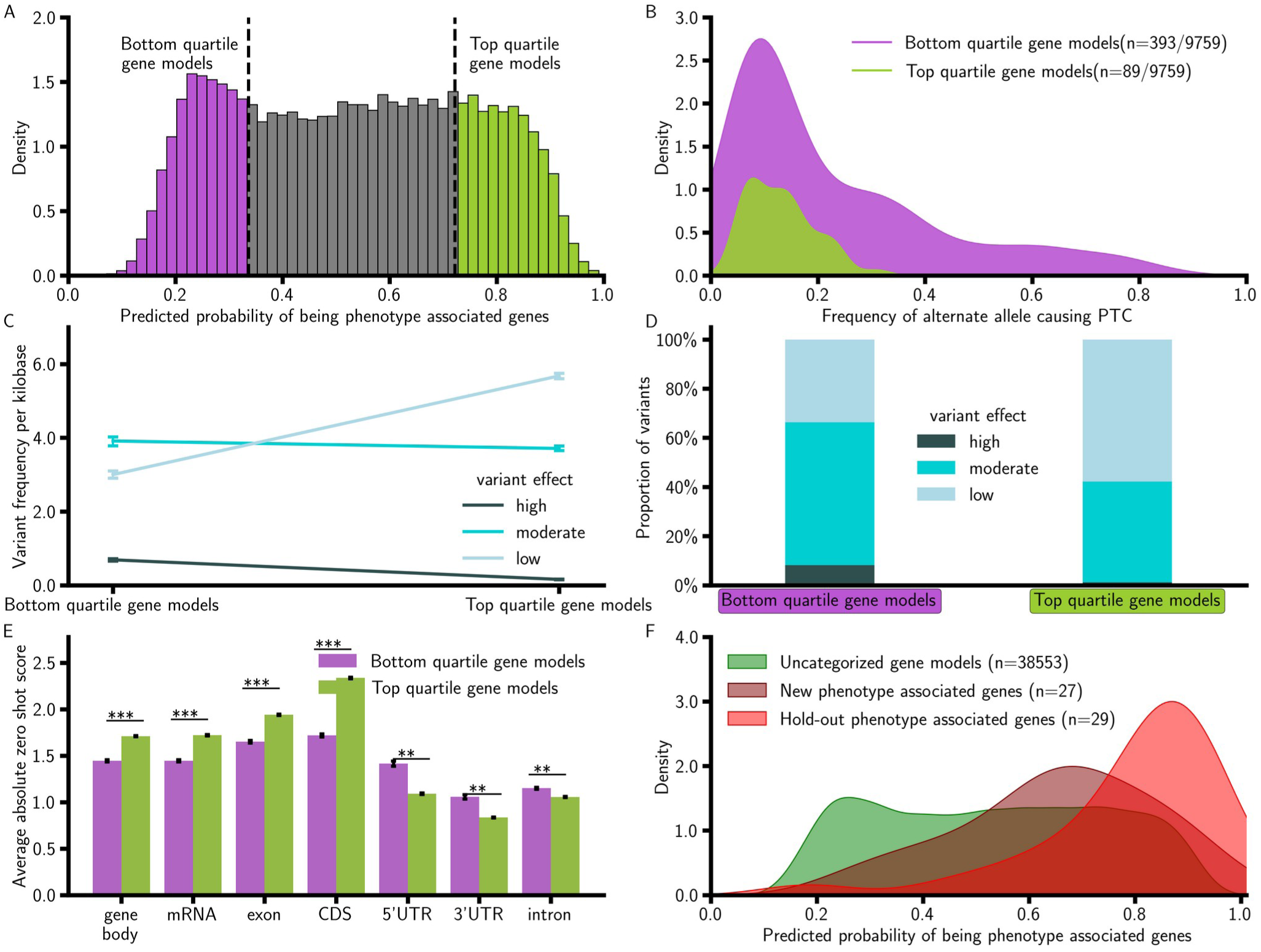
Genes with higher estimated effects on plant phenotypes are conserved in the maize population. A). Bottom and top-quartile gene models were defined from the predicted probability distribution derived from the model trained with the top 100 non-experimental derived features. B). Bottom-quartile gene models show ∼4.6-fold higher odds of containing premature stop codons compared to top-quartile models (*p*−value < 2.26 × 10^−16^, Fisher’s exact test) C). Variant frequency per kilobase in coding sequences of the primary transcript of the gene models, separated by effect type. Bottom-quartile models have higher frequencies of high effect (*p*-value = 2.26e-27) and moderate effect variants (*p*−value = 0), whereas top-quartile models have higher frequencies of low effect variants (*p*−value = 0; two-tailed Mann-Whitney U test). D). Proportional distribution of high, moderate, and low effect variants between quartiles. E). Average absolute zero-shot score for variants estimated from a pre-trained DNA language model, PlantCaduceus, in different parts of the gene models between the bottom-quartile gene models and top-quartile gene models from panel A. ** *p*-value < 1e-20, *** *p*-value < 1e-200 (two-tailed Mann-Whitney U test). F). Predicted probability distribution of newly validated PAGs (*n* = 27, curated from MaizeGDB since November 27, 2021) compared to hold-out PAGs and uncategorized gene models.

Cross-validation and testing on a randomly selected set of hold-out test data are both widely used approaches to estimate the performance of machine learning models, but in many cases, the performance of ML models declines when presented with truly new data relative to subsets of the same dataset used for training (Guan et al., 2025; Lei et al., 2025). A new, curated set of maize PAGs published after our original PAG dataset was finalized was identified from the recent literature (Supplemental Data Set S7). The estimated probabilities assigned to these out-of-sample PAGs remained significantly higher than those of a typical uncharacterized gene (*p*−value = 0.004, two-tailed Mann-Whitney U test, Figure 4F). The average estimated probability assigned to the newly curated set of maize PAG genes was modestly lower than the probabilities assigned to PAGs from the original hold-out test dataset (µ=0.65 vs µ=0.79). However, the estimated probability that a gene in the top 10% of the prediction scores is associated with a phenotype was still approximately 14 times as high as the probability for a gene in the bottom 50% of scores. This ratio was estimated to be 22.5 times using the original hold-out test genes. These results demonstrate that the prediction model retains strong predictive performance even when applied to truly out-of-sample datasets, validating the utility of employing this approach to prioritize candidate genes.

## DISCUSSION

A substantial proportion of gene models remain uncharacterized (Pena-Castillo and Hughes, 2007; Schnable and Freeling, 2011; Stoeger et al., 2018). Importantly, this lack of functional characterization is not limited to individual gene models, as some entire gene families also remain poorly understood. The time and resources required to functionally characterize an individual gene remain costly. As a result, the social structure of research and funding allocation decisions incentivize focusing on well-studied genes. Therefore, more than three-fourths of gene models across humans and plants are understudied and have not been functionally characterized (Schnable, 2020; Stoeger et al., 2018). Although the initial impulse may be to assume that all currently uncharacterized gene models are equally good candidates for characterization, here, we demonstrated that it is possible to train models on relatively simple and generalizable sets of features that dramatically increase the odds of successful reverse genetics characterization. We curated an extended list of phenotype associated genes identified in maize from the 1900s to November 27, 2021 and identified a complementary set of loss-of-function tolerant gene models that can serve as a negative set for model training. All of the genes in this set had evidence of transcription (e.g. expression in RNA-seq studies), many had evidence of translation from proteomic investigations, and more than one-third were conserved at syntenic locations in the sorghum genome, indicating these loss-of-function tolerant genes did not simply represent artifacts produced by *ab initio* gene annotation software. A simple classifier model provided with evolutionary information and genomic sequence-derived features was able to classify gene models in maize into the correct classes with an ROC-AUC of 0.87.

The top quartile of genes predicted to be linked to phenotypes included more than 10% of all maize gene models encoding proteins containing domains of unknown function, more than 300 genes in the top quartile lacked GO annotations, and more than 200 genes lacked any functional description at all, including automated domain annotations. This suggests that our model is not simply learning the properties of well-characterized gene families as uncharacterized genes and members of uncharacterized gene families were not excluded from those predicted to be associated with phenotypes. We were able to assess overfitting or species-specific prediction by testing models trained in maize on known phenotype-associated genes from other species. Models trained in maize generalized well across species, assigning substantially higher predicted phenotype probabilities to genes with known mutant phenotypes in both rice and arabidopsis, two distantly related species with large differences in genome size and gene content.

This ability to generalize highlights the potential of such models to support the annotation of high-confidence gene models in related crops with extensive genomic resources but far fewer cloned and characterized genes such as sorghum (*Sorghum bicolor*) Boyles et al. (2019), as well as orphan or wild species that lack genetic and genomic resources beyond sequenced reference genomes. One potential challenge to employing this approach in species without extensive genomic resources is that one of the features consistently identified as informative for predicting the likelihood of being associated with a phenotype was pan-genome status, i.e., whether a gene has been linked to presence-absence variation. This data type typically requires genomic data (genome assemblies, whole-genome resequencing, comparative genomic hybridization) from multiple individuals of the same species. However, even when we constrained our analysis to core genes found in all characterized individuals, the model retained substantial ability to correctly classify phenotype-associated and loss-of-function-tolerant genes (ROC-AUC=0.61). Similarly, our evaluation of cross-species model performance in rice and arabidopsis was conducted using a model that lacked information on presence-absence variation or pan-genome status. These models achieved their predictive accuracy using only features derived from a single genome assembly, a single set of gene model annotations, and comparisons of syntenic conservation, all features that can be easily generated for any plant with an assembled reference genome.

One potential application of our classifier is predicting the likelihood that disrupting a given gene will be associated with phenotypic changes. Association studies with genetic markers narrow down the genomic region associated with a given trait of interest. There are cases where the causal gene is the nearest one to the associated genetic marker (Sahay et al., 2023), especially when there is a high-density genetic marker across the genome and strong linkage disequilibrium decay (Grzybowski et al., 2023). However, for many plants, markers may be at low density or there may be strong linkage disequilibrium, or in some cases, a combination of both. Having a complementary tool that provides a quantitative score for each gene model indicating the probability that a gene model is linked to a phenotype could guide researchers in prioritizing candidate genes in such association studies. Such a framework could help reduce uncertainty when large genomic intervals contain dozens or even hundreds of annotated gene models. In addition, this tool could provide complementary evidence to traditional QTL mapping or GWAS approaches, ultimately accelerating functional validation. The vast majority of gene models, even in the most studied plants such as maize, rice, and arabidopsis, are yet to be characterized, let alone many orphan crops, which are equally, if not more important (Shrestha et al., 2023). Editing, validating, and characterizing gene models takes a long time and is costly (Stoeger et al., 2018). Quantitative scores for gene models that could be used as confidence scores for prioritizing targets would therefore be invaluable. Such scores could help researchers focus their limited resources on the most promising candidates, thereby accelerating basic discovery and crop improvement. Identifying true genes based on high quantitative scores could mark the starting point for optimizing their functional roles across diverse crop species.

## METHODS

### Definitions of Phenotype-Associated and Loss-of-Function-Tolerant gene sets

The initial set of 272 maize genes whose loss of functional alleles have been experimentally shown to produce a detectable change as an easily observable phenotype was defined by manual curation from a larger set of 603 maize locus records annotated as associated with a phenotype retrieved from the MaizeGDB (Maize Feature Store within the Maize Genomic Database) (Sen et al., 2023) on November 27, 2021. This was done to exclude loci linked to purely molecular phenotypes (e.g., variation in gene expression) or when the evidence for a genotype-phenotype link came from studies using techniques with higher false-positive rates (e.g., association studies). A list of studied rice gene models from the ‘Gene’ tab within the funRiceGenes database was manually curated using the same criteria as for maize gene models until a total of 100 gene models whose loss of function was associated with a detectable phenotype in a publication were identified (Huang et al., 2022). In six cases, a gene identified in the analysis did not correspond to a gene in the sequence-derived feature set calculated from the MSU v7 rice genome annotations, leaving 94 phenotype-associated rice genes. The set of phenotype-associated genes in arabidopsis was retrieved from (Lloyd and Meinke, 2012). Excluding gene models in the Lloyd and Meinke list with phenotypes that were classified as ‘NC; non-confirmed’ resulted in a set of 1,987 phenotype-associated genes for arabidopsis. Out-of-sample maize phenotype-associated genes were identified using a set of 2,582 citations (through April 14, 2025) describing maize loci that had not been associated with a phenotype in our original November 27, 2021 dataset. These citations were derived from 682 unique papers, and screening these publications identified an additional 27 loci that met the criteria for publishing a new genotype-to-phenotype association supported by loss-of-function data. The maize loss-of-function tolerant gene set was defined using a published set of genotype calls derived from whole-genome resequencing of 1,515 maize varieties (Grzybowski et al., 2023). The functional consequence of each of the approximately 46 million biallelic genetic variants (SNPs and InDels) identified within this study were annotated using SNPeff v5.1d (De Baets et al., 2012) and the published gene model annotations of the B73_RefGen_V5 reference genome (Hufford et al., 2021). The primary transcript of each annotated gene model containing a genetic variant that produces a premature stop codon and is present at an alternate allele frequency greater than 0.25 across the population was considered loss-of-function tolerant, resulting in a final dataset of 162 loss-of-function-tolerant maize genes.

### Definition of the feature dataset describing maize gene models

A dataset of 14,418 features describing 39,756 annotated gene models in the B73_RefGen_V5 reference genome was downloaded from the Maize Feature Store in MaizeGDB (Sen et al., 2023). Gene models that were not assigned to any of the ten maize chromosomes (i.e., on unanchored scaffolds) were dropped from the dataset, reducing the total number of gene models to 39,035. Redundant features introduced by the presence of both raw and normalized values in the dataset, features with missing values, as well as features describing tri-peptide amino acid composition, which would otherwise have constituted approximately 75% of the total feature set, were also dropped from the dataset. After excluding these features, 2,639 remained, including features describing gene structure, codon usage, nucleotide and protein composition, and experimentally generated data regarding gene expression atlas, protein expression, chromatin-related features, expression localization, and transcription factor binding sites associated with each gene model in maize. A single non-MaizeGDB-sourced feature, syntenic conservation in sorghum, scored using the dataset published in (Dias et al., 2024), was added, resulting in a final dataset of 2,640 unique features. The final feature set included 31 binary features, 5 categorical features, and 2,604 continuous and discrete features. A total of 350 features were experimentally derived, while the remaining 2,290 features could be directly computed from the maize genome assembly and annotation files.

For use in machine learning applications, categorical features were one-hot encoded, increasing the effective number of variables to 2,675, and continuous and discrete features were min-max normalized across all gene models using the MinMaxScaler function in Scikit-learn v1.3.0 implemented in Python v3.11.4 (Pedregosa et al., 2011).

### Random forest classifier model for prediction

Prior to developing or conducting any analysis, approximately 10% of the gene models were set aside as a separate testing dataset. The data were split into training and testing datasets using a previously proposed family-guided strategy (Washburn et al., 2019) and gene family assignments retrieved from PLAZA (Van Bel et al., 2022) to minimize the risk of data leakage. The testing data set included 29 phenotype-associated genes and 19 loss-of-function-tolerant genes. For each model trained in this study, the remaining 386 labeled gene models were used to optimize the hyperparameters and train the final models. The number of trees in a model, number of features for splitting each tree, maximum depth of the tree, minimum number of samples required to split a node, minimum number of samples required to be at a leaf node, and whether bootstrap samples were used when building trees were optimized via three-fold cross-validation of the training dataset using the RandomizedSearchCV function in Scikit-learn v1.3.0 implemented in Python (Pedregosa et al., 2011). The optimal set of hyperparameters identified through this process for each model was then used to train the respective final model using the whole training data. ROC curves were plotted based on the model’s performance on the hold-out test dataset, and the AUC was used to evaluate the model’s performance.

Feature importance scores for both models using 2,675 and 2,290 features were calculated using the built-in feature_importance_ function of a trained random forest model from the Scikit-learn library implemented in Python. This method estimates the importance of each feature based on the mean decrease in impurity across all trees in the ensemble. We were only able to generate values for 95 of the 100 maize features of greatest estimated importance in rice and arabidopsis. The five remaining features (Core Gene, Dispensable Gene, gene age, PseDNC Xc1 TT, and Stacking Energy) were not included in cross-species prediction. A new model trained on maize data using only those 95 genomic features and the same procedure described above, was employed for cross-species predictions.

### Validation of model predictions

Variant density was calculated by dividing the number of variants from the same resequencing-based dataset used to identify loss-of-function-tolerant genes occurring in the CDS of the primary transcript by the CDS length of the primary transcript of that same gene. Additional density values were generated by classifying genetic variants present within the primary transcript of a given gene into low effect, moderate effect, modifier effect, and high effect categories via SNPeff (De Baets et al., 2012). The functional impact of genetic markers present between the annotated start and stop sites of each gene of interest was estimated using zero-shot scores provided by PlantCaduceus (Zhai et al., 2025). The absolute value of the log-likelihood difference between reference and alternate alleles was used when the reference or the alternate allele was assigned low relative probabilities, both of which indicate the presence of functional differences segregating in the population. InterProScan V5.75 was used to identify gene models whose translated CDS coded for one or more domains of unknown function (Jones et al., 2014). Tests for GO enrichment or purification were performed using the Python library Goatools v1.4.12 (Klopfenstein et al., 2018). The Gene Ontology annotation file, “Zm00001eb.1.fulldata.txt.gz”, for B73 reference genome version 5 was downloaded from maizeGDB (Hufford et al., 2021). To exclude bias introduced by the presence of greater proportions of some gene sets representing entirely uncharacterized proteins, the background set of all analyses was defined as the set of genes associated with one or more GO terms unless otherwise specified. GO term enrichment analysis was calculated as the fold change between the proportion of gene models annotated with a given GO term in the study set and its proportion in the background population set of the gene models.

### Curation of rice and arabidopsis gene model features

The rice genome was retrieved from the Rice Genome Annotation Project (Hamilton et al., 2025; Kawahara et al., 2013), and the arabidopsis genome was retrieved from The Arabidopsis Information Resource (TAIR) (Lamesch et al., 2012). For both species, only data from primary transcripts were used, and genes annotated as transposable elements (TEs) were excluded. Length features including 3’ UTR length, 5’ UTR length, total exon length, and average exon length were directly calculated from gene annotations in the GFF3 file. K-mer ratios were derived using a custom R script that integrated the FASTA sequence with GFF3 annotations. Codon usage frequency, relative synonymous codon usage (RSCU), and GC content at the third codon position (GC3s) were also computed from representative CDS.

Multiple protein sequence descriptors were extracted from one representative peptide sequence per gene using the protr R package (Xiao et al., 2015). After removing stop codons and filtering out sequences shorter than three amino acids, the following descriptor types were computed: triad composition, Geary, Moran, and Moreau-Broto autocorrelations, dipeptide composition, pseudo-amino acid composition (PAAC), quasi-sequence order (QSO), and composition-transition-distribution (CTD: CTDC, CTDD, CTDT). Custom functions with built-in error handling were applied to extract specific subsets of features.

All computations were implemented in parallel to accelerate processing. Syntenic rice genes were identified based on orthology with sorghum genes and directly obtained from the supplementary materials of a previously published study (Dias et al., 2024). Syntenic arabidopsis genes were inferred via comparison with *Brassica rapa* genes using the latest reference genome and the SynOrths pipeline (Cheng et al., 2012). Features such as core label Core Gene, core label Dispensable Gene, gene age, PseDNC Xc1 TT, and StackingEnergy were not included, as no reliable public resources were found, and they could not be readily derived from sequence or annotation data using straightforward computational methods.

## Acknowledgements

The authors would like to thank scientists from MaizeGDB, especially Dr. Maggie Woodhouse and Dr. Eddy Cannon, for providing information, access to the data used in this study, and help with curating the new phenotype-associated gene models. The authors would also like to thank Jingjing Zhai for providing access to zero-shot scores for maize variants published in MaizeGDB. This research was supported in part by the U.S. Department of Agriculture, National Institute of Food and Agriculture, under award numbers 2020-67021-31528 and 2020-68013-30934, and the AI Research Institutes program supported by NSF and USDA-NIFA under the AI Institute for Resilient Agriculture, Award No. 2021-67021-35329. Zhongjie Ji was additionally supported by a Heuermann Postdoctoral Fellow award from the University of Nebraska–Lincoln.

## Author contributions statement

J.C.S. conceived of the project. N.S. processed and analyzed the data. Z.J. generated data for validation of the model on rice and arabidopsis. J.C.S. provided supervision, guidance, and feedback on the design of analyses and approaches. N.S. drafted the paper with input from J.C.S. All authors contributed to the revision of the final manuscript.

## Conflict of Interest

James C. Schnable has equity interests in Data2Bio, LLC; and Dryland Genetics LLC and has performed paid work for Alphabet. He is a member of the scientific advisory board Aflo Sensors. The authors declare no other competing interests.

## Supplemental Data

- **Supplemental Data Set S1.** List of 272 phenotype-associated gene models validated through loss-of-function experiments, including corresponding publications or database links.
- **Supplemental Data Set S2.** Test statistics comparing phenotype-associated gene models and loss-of-function–tolerant gene models across the 2,604 genomic, structural, evolutionary, and sequence-derived features used in this study.
- **Supplemental Data Set S3.** Predicted probability of each maize gene model being phenotype-associated, generated using the top-100-feature random forest classifier.
- **Supplemental Data Set S4.** Top 100 features identified by the random forest classifier, ranked by Gini impurity–based importance.
- **Supplemental Data Set S5.** Predicted probabilities of phenotype association for rice and Arabidopsis gene models using the maize-trained classifier.
- **Supplemental Data Set S6.** Gene Ontology enrichment results for maize gene models in the top and bottom quartiles of predicted phenotype-association probability.
- **Supplemental Data Set S7.** Predicted phenotype-association probabilities for 27 newly characterized maize phenotype-associated genes.

## Notes

### Summary of Updates

We made changes to the abstract section. The rest of the section remains the same along with data and analysis.

